# On the evolution of dispersal strategies under the costs of acquisition of private and social information

**DOI:** 10.1101/2024.12.17.628840

**Authors:** Antoine Sion, Matteo Marcantonio, Stefano Masier, Elio Tuci

## Abstract

Dispersal between patches of suitable habitat is a key behaviour for the survival of animal populations and is impacted by rapid environmental changes. Animals must cope with several costs related to the dispersal process, such as expenses in energy for acquiring information or increased mortality risks. There is a growing interest in studying how the costs of acquisition of information influence the dispersal behaviour of species, and particularly how the use of social and private information can shape this behaviour over multiple generations. Current models of dispersal rarely incorporate both sources of information, and there is specifically a lack of modelling studies taking into account costs of acquisition for private and social information. We develop an agent-based model simulating a population of butterflies with genetic factors linked to the acquisition of both types of information and their associated reproductive costs. We show that different costs, environmental variability conditions and sensory abilities result in various dispersal behaviours and have an impact on the fitness of the population. In stable environments, low-cost information is used by a varying proportion of agents to disperse, but when the cost rises lightly, all agents stay uninformed. In highly variable environments, the same trend is observed, but agents rely on information even if the cost of acquisition increases up to twice compared to stable environments as it provides an evolutionary advantage. Agents with a limited perceptual range use both information sources equally in variable environments, and those with a bigger perceptual range rely exclusively on private information to take dispersal decisions, except at free cost of acquisition. Globally, the use of information induces a higher fitness for the population in stable environments if costs of acquisition are free or very low. In variable environmental conditions, the highest fitness is maintained with a limited perceptual range when the total cost rises up to twice the value found for stable environments. With a bigger perceptual range, the highest fitness is maintained for the whole range of total costs studied.

## 1 Introduction

Human activities are rapidly altering the environment, impacting animal mobility and habitats. This phenomenon, known as Human-Induced Rapid Environmental Changes, imposes behavioural adaptations among animals [62]. There is a growing research interest in how animals adapt to these changes, with their responses playing a vital role in either exacerbating or mitigating environmental pressures [6, 46, 56]. The study of these adaptations is increasingly recognised as essential for understanding evolutionary responses to environmental changes, with empirical and theoretical evidence highlighting its significance [6, 51, 16, 53, 15].

One key behaviour affected by environmental changes is dispersal between patches of suitable habitat [49, 9, 28]. This type of movement underpins population survival as it provides important benefits such as escaping competition, avoiding inbreeding and increasing the quantity of exploitable resources. Nevertheless, it is also associated with significant energetic costs and increased mortality risks [58, 5, 11, 27]. The chain of decision-making that brings to dispersal is complex, influenced by an animal’s current state, its ability to move and navigate, and the environment [32, 3]. The process of evaluating the quality of the environment, for example, in terms of abundance of resources or with respect to the presence of predators, is generally based on the integration of different sensory pathways, and it is strictly associated to the sensory machinery of species [54]. Such information is often highly accurate, but it is not always available to organisms or has a higher acquisition cost (refer to box **Box 1** for definitions). Another more widely available source of information that is recognised as critical for species facing problems related to Human-Induced Rapid Environmental Changes is “social information”, that is, the use of information provided by the observations of other individuals behaviour. Recent evidences show how social information is used also by non-social species, particularly in the context of resource availability [11, 60, 7, 33]. Such type of information is pervasive in the environment and provides organisms with additional cues to localise high quality resources, such as food or oviposition sites, at low energetic cost [13, 55]. Organisms tend to rely more on social information when the frequency of change of environmental conditions makes individual knowledge relatively inaccurate [22, 52, 34]. Among the most simple and widespread “social information” that can be acquired by organisms, is the density of conspecifics, which has been found affecting the dispersal of multiple taxa, such as insects [see 35, 2, 17, 20]. Principally, the presence of conspecifics is considered a widespread cue to evaluate habitat quality across taxa [e.g., 29, 14], and over 80% of studies on this topic have found that social cues have an attractive effect [8]. However, crowding is also associated with smaller size, lower fecundity, reduced offspring viability, and higher rates of offspring starvation in various invertebrate and vertebrate taxa [61, 57, 38].

When exploring how animals, such as butterflies, gather and use information to make decisions, it is essential to consider not just the type of information (i.e., private or social) but also where this information comes from. This can include their current patch of habitat, or neighbouring areas, whose extension varies depending on species specific perceptual capabilities. In many species, this perceptual range is defined by how well they can integrate sensory inputs, such as visual and olfactory cues, and can be assumed to decrease as an inverse function of distance. In the case of butterflies, their sensory capabilities allow them to perceive their habitat from a few meters up to about 100 meters [23, 49].

The availability, the acquisition and the integration of different kinds of information allow organisms to maximise dispersal decisions, and by extension, their fitness. Despite its significance, experimental research into how environmental cues and social information interact to shape population dynamics and movement behaviour in varying habitats is limited. This lack of study is largely due to the challenges of accounting for multiple information sources and for environmental variability [33]. Simulation models that mimic the life cycles and evolution of individual organisms offer opportunities to delve into this issue [5]. They provide a controlled setting that allow researchers to explore how the collection of information, its associated costs, environmental changes and movement choices interact.

Simulation models implementing dispersal tend to only consider the use of conspecific density as a source of information that agents exploit to decide whether to disperse or not [see 47, 40, 63, 1, for example]. Recent studies using simulation models have integrated socially derived and individually gathered information to investigate how multiple sources of knowledge bear upon the results of decision-making processes related to dispersal. Ponchon et al. [45] showed that dispersal decisions relying on private and social information allowed populations persistence in spatially variable environments, whereas random dispersal led them to extinction. The same type of model is later applied to study species range expansions [44], showing that expansions are slower when emigration decisions are informed by assessment on habitat quality. Enfjäll and Leimar [18] compared scenarios with increasing degrees of available social and private information, reporting that the proportion of dispersing agents progressively decreases with increasing overall information [as also found in 24, 41]. However, this model did not consider the cost of information acquisition on the reproductive success of the agents. Bocedi et al. [4] included a reproductive cost of information acquisition. However, their contradictory results on density-dependent dispersal have been criticised due to a series of distortions introduced into their modelling framework [see 43]. In particular, Poethke et al. [42] highlighted how the cost of information acquisition influences the dispersal behaviour of populations solely based on density-dependent information. They found that high costs deter agents from utilising this information in dispersal decisions in environments where habitat quality varies greatly. In scenarios of increasing environmental variability, the reliance on information acquisition for dispersal increases if the cost remains constant. Notably, there is a lack of a modelling study that incorporates informed dispersal based on both private and social information, factoring in their total acquisition costs. Our study aims to address this gap by examining the total acquisition costs associated with social and private information in dispersal decisions.

We develop an agent-based model that simulates the population dynamics of butterflies. This model enhances and expands the framework proposed by [18] incorporating genetic factors that drive the evolution of social and private information gathering. As energy available to organisms is often limited, investing in the sensory machinery underpinning information acquisition and integration comes with a trade-off with other traits, such as fecundity [5]. We have therefore added reproductive costs associated with acquiring information to the model. We study the evolutionary trajectory of genes associated with the acquisition of environmental and social information and we show how this influences population dynamics and the choice of moving between patches of habitat. We chose butterflies as our focus taxon because they are widely used as a model organism to understand population dynamics and movement. Moreover, butterflies are crucial indicators of environmental health and play an important role in highlighting the current biodiversity crisis [59, 27].

Our findings reveal that the interplay between environmental variability and the cost of gathering information shapes the evolution of strategies within butterfly populations. In stable settings, agents that depend on direct, private observations are more likely to succeed if information has very low fitness costs or when it is free. Scenarios with increased cost of information select for agents uninformed on the environmental and the conspecific density of their local or neighbour habitat patch. Conversely, in highly variable environments, populations that adapt by using a mix of social and non-social cues thrive, even when obtaining this information is costly. In these highly variable environments, there is a clear advantage for agents capable of interpreting surrounding cues of non-social signals. Our results demonstrate how adaptability to a wide range of environmental cues is crucial for survival under Human-Induced Rapid Environmental Changes.

## 2 Material and methods

Our model simulates the life cycle, dispersal and evolution of a population of butterfly agents in a finite arena composed of squared habitat patches (hereafter, also referred to as cells). In this section, we explain the structure of the model concisely, whereas a complete model description is reported in appendix A.

At the beginning of each model simulation, 40,000 agents (sex ratio 1:1) are randomly placed in a grid of 20 × 20 cells forming a torus. Each patch has a carrying capacity randomly drawn from a normal distribution at each new generation. Only females are able to disperse into adjacent patches once per generation. Dispersal decisions are evaluated by first acquiring private or social information. Private information refers to the knowledge of the patch carrying capacity, while social information refers to the number of conspecifics on a patch. We explore two scenarios. In one scenario (hereafter, referred to as RPR), agents have a restricted perceptual range, which make them capable of acquiring information from the natal patch only. In the other scenario (hereafter, referred to as EPR), agents have an extended perceptual range, which make them capable of acquiring information from the natal patch and from an additional randomly selected adjacent patch. In RPR, two binary genes, one for social (*s*_1_) and one for private information (s_2_), determine whether or not an agent uses private and/or social information about the natal patch to compute its probability to disperse. When any of these two genes are active, the contribution that each type of exploitable information brings to the computation of the agent’s probability to disperse is determined by two additional real-valued genes, one associated to private (*a*_1_) and one to social information (*b*_1_). In particular, each real-valued gene determines whether private/social information contributes to increase or decrease the agent’s probability to disperse and also the magnitude of this contribution. In EPR, whether private and/or social information are exploited and their respective contribution is regulated by six genes, two binary genes for the exploitation of social/private information (*s*_1_ and *s*_2_), two real-valued genes for the contribution of the carrying capacities of the two patches (*a*_1_ for the natal, and *a*_2_ for the randomly selected adjacent patch), and two real-valued genes for the contribution of the numbers of conspecifics of the two patches (*b*_1_ for the natal, and *b*_2_ for the randomly selected adjacent patch). In both scenarios, the genes are passed asexually to the next generation and mutate at each generation.

### Box 1

#### Definitions

- Information: Anything that reduces uncertainty affecting the unfolding of a decision-making process, for example agents dispersal [13, 12, 48].
- Private information: Information about the quality of the habitat available to agents through direct observations of the physical world, i.e., the carrying capacity of a patch of habitat [36].
- Social information: Information about the quality of the habitat that is indirectly available to individuals by monitoring others’ interaction with the surrounding environment, i.e., density of conspecific in a habitat patch.
- Cost of information: Negative impact on the reproductive success of female agents linked to the process of (any) information acquisition.
- Free information: Information that, when acquired, does not negatively impact the reproductive success of the female agents obtaining it.

The exact sequence of events occurring at each generation is shown in the diagram in Figure 1. At the beginning of a simulation (i.e., at generation 1), agents’ genes and patches carrying capacities are initialised by sampling values from their respective distributions. Then, during the reproduction phase, the local number of agents for the next generation is computed for each patch from the number of females and the patch quality. For each new agent, its genes are chosen in the local pool of mated females at random following a weighted distribution with replacement. Each female has a reproductive weight associated to information acquisition. Generations are non-overlapping with old agents dying before the birth of the new ones. Environmental variation is introduced by resampling the quality of each patch from a normal distribution. Females mate in the presence of males in their local patch. Finally, females take a dispersal decision by computing their probability of dispersal, taking into account the type of information enabled by their genes. Fully informed females have the smallest reproductive weight whereas uninformed females have the highest.

**Figure 1:**
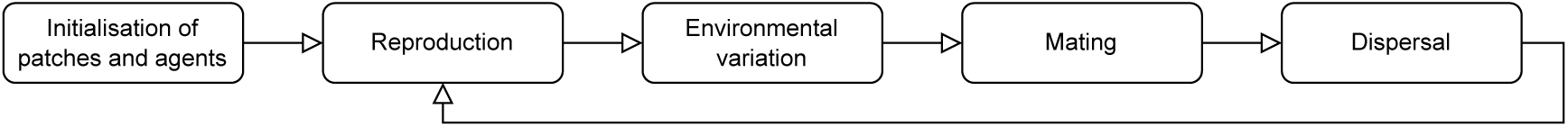
Sequence of events in our model.

The simulation model investigates how the frequency of the genes above-mentioned changes in multiple experimental conditions, given by all the possible combinations of two scenarios (RPR and EPR illustrated above) and three types of environmental state characterised by the parameter *σ* regulating how the quality of patches varies between following generations (i.e., very slightly with *σ* = 1, slightly with *σ* = 10, largely with *σ* = 100). The resulting six experimental conditions are referred to as:

- Condition A (Cond. A): scenario RPR and *σ* = 1;
- Condition B (Cond. B): scenario EPR and *σ* = 1;
- Condition C (Cond. C): scenario RPR and *σ* = 10;
- Condition D (Cond. D): scenario EPR and *σ* = 10;
- Condition E (Cond. E): scenario RPR and *σ* = 100;
- Condition F (Cond. F): scenario EPR and *σ* = 100;

For each condition, we vary the cost of exploiting private and/or social information on agent’s fertility regulated by the parameter *r*_min_ ∈ {0.75, 0.8, 0.85, 0.9, 0.95, 1.0}. The lower the value of *r*_min_, the higher the costs on fertility for exploiting private and/or social information. When *r*_min_ = 1, these costs are zero, meaning that the acquisition of information is free for all agents. Note that, we assume that the reproductive costs of acquiring private and social information are equal, but this assumption can be easily modified in future works. For each set of parameters (i.e., for each condition), we run 20 randomly seeded simulations of our model, with each simulation lasting 20,000 generations.

## 3 Results

In this section, we illustrate the results of our study, by first showing, for each condition, and for different costs (i.e., *r*_min_ ∈ [0.75, 1.0] ), the distributions of the population size (see Figure 2) at the last generations (i.e., at generation 20,0000). Note that, the between generation variability in the number of agents within each single run is very minimal when attaining the evolutionary steady state with standard deviations values being around 50 for *σ* = 1, 200 for *σ* = 10 and 1500 for *σ* = 100 (data from the last 100 generations).

**Figure 2:**
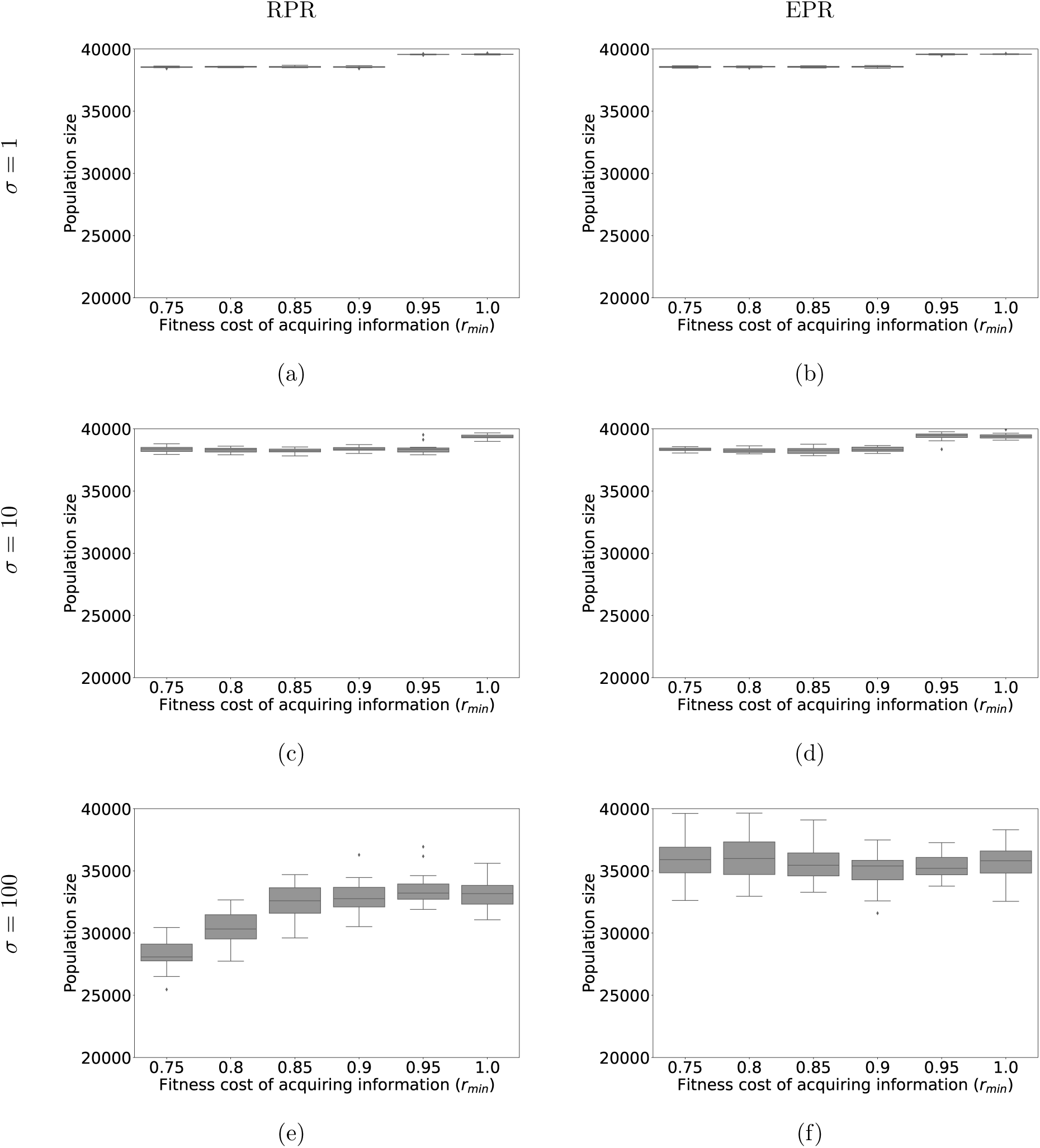
Graphs showing the distribution of population sizes at generation 20, 000 for different values of *r*_min_. Figures (a), (c) and (e) refer to conditions in which the agents can access only information concerning the natal patch with *σ* = 1 in (a), *σ* = 10 in (c), and *σ* = 100 in (e). Figures (b), (d) and (f) refer to conditions in which the agents can access information concerning the natal and a randomly selected adjacent patch with *σ* = 1 in (b), *σ* = 10 in (d), and *σ* = 100 in (f).

When environmental variability is very low (*σ* = 1, see Figure 2a for RPR, and 2b for EPR), or low (*σ* = 10, see Figure 2c for RPR, and 2d for EPR), final generation population sizes are within [37, 500, 40, 000] agents with relatively small variability i) between different runs given *r_min_*; and ii) between different costs (i.e., for different values of *r_min_*). Moreover, the same very mild decrement in the population size for progressively lower values of *r_min_* is observed in both scenarios (see Figure 2a and 2c for RPR, and Figure 2b and 2d for EPR). When environmental variability is high (*σ* = 100, see Figure 2e for RPR, and 2f for EPR), the population size tends to stabilise at lower values than those observed for *σ* = 1 and *σ* = 10, with final populations of the RPR scenario having smaller sizes than those of the EPR. While in RPR the smaller *r_min_* the smaller the population size (see Figure 2e), in EPR this trend does not emerge (see Figure 2f). Statistics on the population size suggest that, when the environment varies very little or little, the population size tends to stabilise on relatively close values for different fitness costs for exploiting private and/or social information. When the environmental state is highly variable, the population sizes tend to be smaller than those observed in stable environmental states, and with larger between runs variability.

Figure 3 shows, for each condition, and for different costs (i.e., *r_min_* ∈ [0.75, 1.0]), the distributions of the frequency of allele *s_1_* = 1 (see Figure 3, white boxes for social information) and *s_2_* = 1 (see Figure 3, grey boxes for private information) at the last generations (i.e., at generation 20,0000). As for the population size, the between generation variability in the frequency of both alleles within each single run is very minimal when attaining the evolutionary steady state, with standard deviations values being around 0.02 in all environmental conditions (data from the last 100 generations). When private and social information can be acquired at no fitness cost, and the environment is highly variable (*σ* = 100), in scenario RPR both alleles are fixed (see Figure 3e, white and grey boxes for *r_min_* = 1); in scenario EPR, while *s_2_* = 1 is fixed (see Figure 3f, white box for *r_min_* = 1), *s_1_* = 1 tends to appear in slightly more than half of the population. When progressively higher costs are associated to private and social information (i.e., for *r_min_ <* 1), both alleles tend to reduce their frequency, but the nature of this trend largely differs with respect to the experimental condition. When the environment varies very little (*σ* = 1) or little (*σ* = 10), both alleles disappear from the population for *r_min_ <* 0.95. That is, a small fitness cost for accessing private and social information is enough to make these strategies extinguished (see Figure 3c for RPR, and 3d for EPR). When the environment is highly variable (*σ* = 100), the progressive increase of fitness costs makes both alleles progressively disappear from the population when *r_min_ <* 0.8 in scenario RPR (see Figure 3e). On the contrary, in scenario EPR, while the allele for accessing social information extinguishes as soon as fitness costs appear (see Figure 3f, white boxes for *r_min_ <* 1.0), the allele for accessing private information tends to remain either fixed or with a extremely high frequency in the population (see Figure 3f, grey boxes, for *r_min_ <* 1.0).

**Figure 3:**
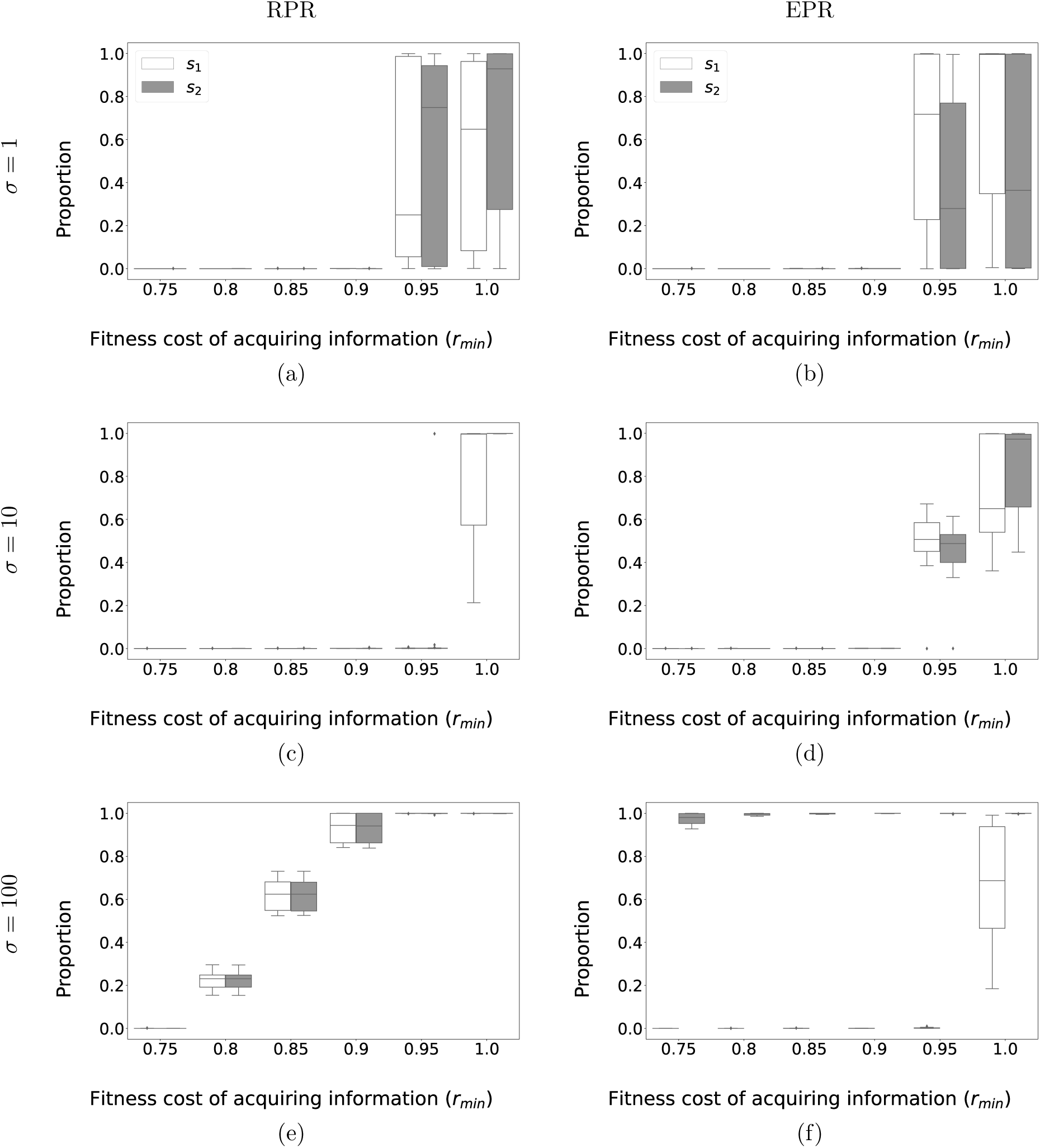
Graphs showing the distribution of frequency of allele *s*_1_ = 1 (white boxes) and *s*_2_ = 1 (grey boxes) at generation 20, 000 for different values of *r*_min_. Figures (a), (c) and (e) refer to conditions in which the agents can access only information concerning the natal patch with *σ* = 1 in (a), *σ* = 10 in (c), and *σ* = 100 in (e). Figures (b), (d) and (f) refer to conditions in which the agents can access information concerning the natal and a randomly selected adjacent patch with *σ* = 1 in (b), *σ* = 10 in (d), and *σ* = 100 in (f).

In those cases, in which the allele *s*_1_ = 0 (social information) and/or *s*_2_ = 0 (private information) are fixed (see, for example, Figure 3a, for *r_min_ <* 0.95), the other genes (i.e., the genes *a*_1_ and *b*_1_ in scenario RPR, and genes *a*_1_, *b*_1_, *a*_2_ and *b*_2_ in scenario EPR) drift without contributing to the agents’ fitness. In all the other cases, the values of genes *a*_1_, *b*_1_ for scenario RPR (see Figure 4a, 4c, 4e), and genes *a*_1_, *b*_1_, *a*_2_ and *b*_2_ for scenario EPR (see Figure 4b, 4d, 4f), determine how private and social information is exploited by the agents to compute the probability of dispersal. In the RPR scenario, when the environment varies very little (see Figure 4a, *σ* = 1), both genes *a*_1_ and *b*_1_ only slightly contribute to reduce the probability of dispersal. For progressively more variable environmental conditions (see Figure 4c for *σ* = 10, and 4e for *σ* = 100) gene *a*_1_ (i.e., for private information) contributes to reduce the probability of dispersal, while gene *b*_1_ (i.e., for social information) contributes to increase the probability of dispersal. Thus, the probability of female dispersal decreases with increasing quality of the natal patch or decreasing conspecific density.

**Figure 4:**
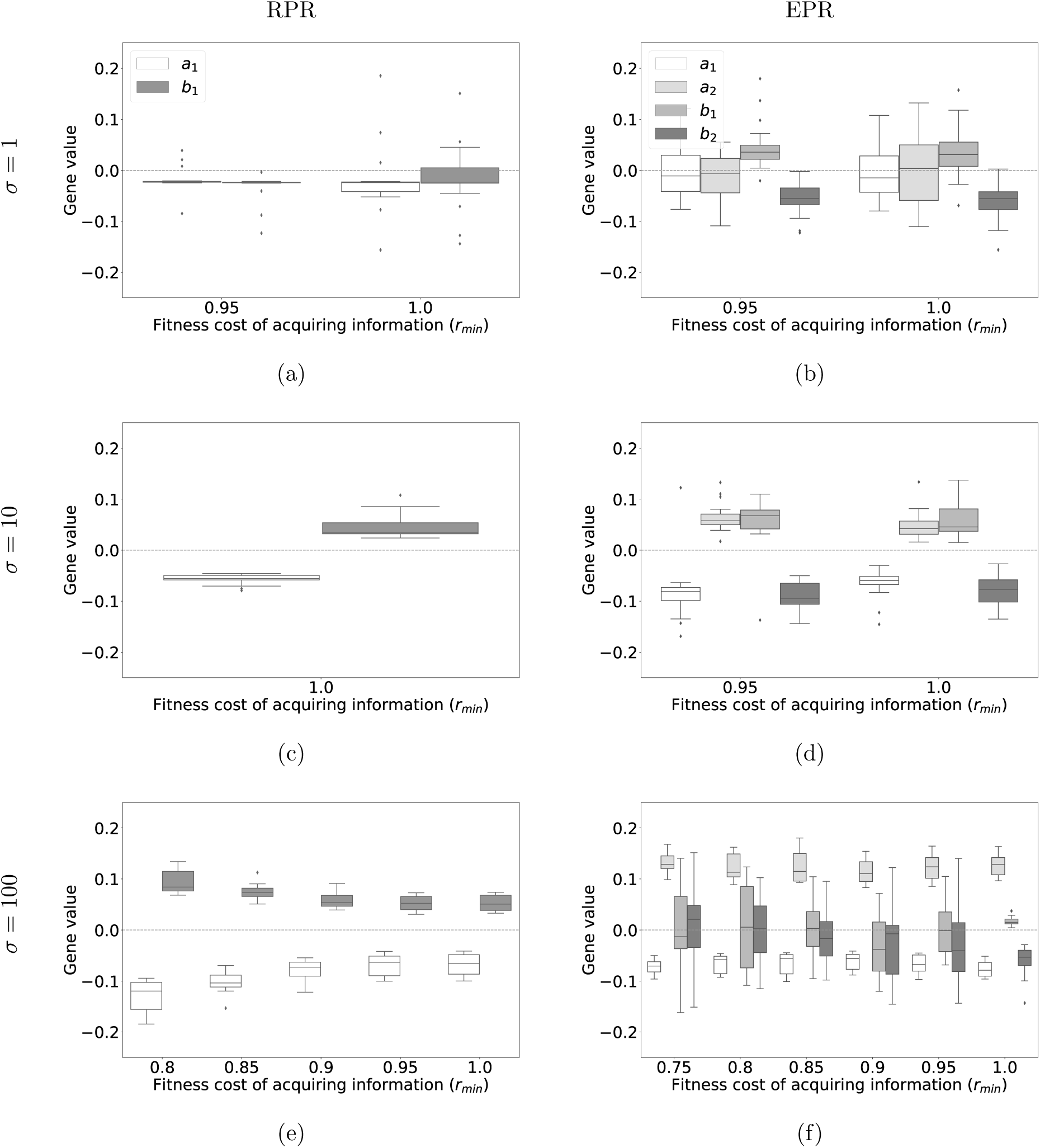
Graphs in figures (a), (c) and (e) show the distribution of genes *a*_1_ (white boxes) and *b*_1_ (grey boxes) in scenario RPR, for *σ* = 1, *σ* = 10 and *σ* = 100, respectively. Graphs in figures (b), (d) and (f) show the distribution of genes *a*_1_ (white boxes), *a*_2_ (light grey boxes), *b*_1_ (grey boxes), and *b*_2_ (dark grey boxes) in scenario EPR, for *σ* = 1, *σ* = 10 and *σ* = 100, respectively. In each graph, the costs defined by *r*_min_ are those in which the allele *s*_1_ = 1 and/or *s*_2_ = 1 are present in the final generation populations.

In the EPR scenario, when the environment varies very little (see Figure 4b, for *σ* = 1), genes *a*_1_ and *a*_2_ do not appear to contribute significantly to the dispersal strategy while *b*_1_ slightly contributes to increase the probability of dispersal and *b*_2_ tend to slightly contribute to reduce it (see Figure 4b, grey boxes). When the environment varies little (see Figure 4d, for *σ* = 10), genes *a*_1_ (see Figure 4d, white boxes) and *b*_2_ (see Figure 4d, dark grey boxes) contribute to reduce the probability of dispersal, while gene *a*_2_ (see Figure 4d, light grey boxes) and *b*_1_ (see Figure 4d, grey boxes) contribute to increase the probability of dispersal. When the environment is highly variable (see Figure 4f, for *σ* = 100) gene *a*_1_ contributes to decrease the probability of dispersal (see Figure 4f, white boxes) while gene *a*_2_ contributes to increase the probability (see Figure 4f, light grey boxes). The other genes tend to have either very little effect or to reduce the probability of dispersal. Overall, for moderate to high environmental variations, females are more likely to disperse towards high quality patches populated with a low number of conspecifics, especially if the natal patch has poor quality and/or is crowded.

To summarise, our results show that highly variable environmental conditions lower population sizes because such environments correspond to harsh foraging conditions, directly affecting reproduction. The propensity to use private and social information tends to disappear when costs associated to the use of these information sources increase, but the way in which populations stop using information depends on the environmental variability. The higher the variability, the longer it takes for these genes to become extinct. The gene for using private information never disappears in highly variable environments, which means that it is more evolutionary efficient to pay the costs of testing the quality of a patch, rather than not paying these costs and running the risk of ending up in low quality patches. When using information in the RPR scenario for highly variable environments, females disperse away from low-quality natal patches with high population densities. In the EPR scenario with access to information from an adjacent patch, harsh environments select for behavioural strategies in which females aim to disperse to sparsely populated, high quality patches.

## 4 Discussion

The unprecedented pace of Human-Induced Rapid Environmental Changes ([50]) presents environments changing more rapidly than species can adapt during their evolutionary history [37]. This rapid change challenges the long-term viability of species, making it imperative for them to dynamically adjust to their habitats within these altered ecosystems to survive. In such scenario, the ability of organisms to gather and process accurate environmental information to take informed dispersal decision is pivotal.

### 4.1 Perceptual range, environmental variation and population size

Our findings suggest that in variable environments, agents with an extended perceptual range have a higher fitness and therefore reach a higher population size than agents with a more restricted perceptual range (e.g., the natal patch). Several empirical evidences suggest that species with a limited perceptual range may be impacted more by habitat fragmentation than species with an extended perceptual range [30]. Theoretical studies have confirmed this relationship by exploring how the variation in perceptual range changes the fitness of a given population. For example, [19] developed a continuous time-space model to study the optimal perceptual ranges for foragers in dynamic landscapes, showing that non-local information is highly valuable especially in dynamic environments. Similarly, results in [39] highlighted that a higher perceptual range enhances connectivity and the response from animals to habitat fragmentation.

In more stable environments, the importance of an extended perceptual range diminishes and no difference between population sizes can be found than populations with a restricted perceptual range. It is well-known that individuals belonging to the same species employ different dispersal strategies depending on the level of habitat variability they belong to [31]. However, it is unclear if the phenotype is selected by the environment or if individuals with a certain phenotype thrive in more or less variable habitats due to intraspecific competition. Our results suggest the former of the two alternative hypotheses since agents with specific dispersal phenotypes were selected under high environmental variability, without our model inherently integrating direct intraspecific competition. Moreover, this result show that relying on diverse sources of information can be a valuable evolutionary advantage to adapt to high environmental variations, but it is irrelevant in stable environments.

### 4.2 The cost of information on fitness and dispersal

If, on the one hand, an extended perceptual range may allow for the acquisition of more information to underpin dispersal decision, on the other hand, this information comes with a trade-off with other traits, such as fecundity [5].

In stable environments, information may be regarded as of little value to organisms and thus not worth the energetic investment. An organism dispersing into such environments has a high likelihood to land onto a patch with characteristics similar to its natal patch. Results from our simulations showed that when information is free or associated with a very low fitness cost, agents exploit it to take dispersal decisions. Thus, informed movements can be more advantageous than random movements also when habitat patches have similar qualities. But, as the cost of information increases, agents rapidly abandon its use.

In environments with higher variability, organisms gain an evolutionary advantage by acquiring information, even if doing so incurs higher costs. For example, butterflies dispersing in highly variable environments, such as marginal habitat surrounded by inhospitable agricultural areas, often need to leave their natal patch to locate new less crowded habitat conducive to their survival and reproduction. However, this dispersal process is fraught with risks, including a high likelihood of death [49]. Similarly, common toads exhibit different dispersal behaviours depending on the fragmentation level of the area they originated from [25], and rainforest birds were more or less adept in crossing stretches of unsuitable matrix depending on their population of origin [10]. In these situations, the acquisition of information becomes crucial. It ensures that decisions to disperse are made only when there is a definitive benefit in terms of habitat quality improvement. This higher investment in information acquisition is similar to the results reported in [42], with agents able to exploit more accurate information selected in highly variable environments.

### 4.3 Information availability underpins emerging dispersal strategies

In this analysis, we observed that agents’ dispersal strategies, influenced by their perceptual range and environmental conditions, are crucial for optimising fitness. In stable environments, agents with limited perceptual capabilities predominantly utilise private information for dispersal, only rarely using social information (e.g., population abundances). This trend changes in variable environments, where reliance on social information increases, indicating a balanced use of both information types for optimal dispersal decisions. Notably, agents with broader sensory capacities utilise both information types in stable settings, showing a preference for social cues in highly stable scenarios. Conversely, in more variable environments, butterfly agents primarily depend on private information, suggesting that direct knowledge of habitat quality may be enough to take dispersal decisions without the need of social cues. This pattern was suggested on the assumption of equal costs for acquiring both types of information. Should the cost differ (see for example [26]), we anticipate the evolution of dispersal behaviours that favour the more economically acquired information [33].

The study of gene evolution in our model offers some insights into the behaviour of agents, showing consistent trends across scenarios where information influences dispersal. High conspecific numbers in the natal patch and lower numbers in adjacent parches, along with comparative habitat quality, are key determinants of female dispersal probability. This detailed examination suggests the adaptive strategies agents employ in response to environmental cues and perceptual limitations, highlighting the complex interplay between genetics, behaviour and ecological dynamics.

## 5 Authors’ contributions

**Antoine Sion:** Conceptualisation, Methodology, Software, Validation, Investigation, Writing - original draft, Visualisation. **Matteo Marcantonio:** Conceptualisation, Methodology, Writing - original draft, Writing - review and editing, Supervision. **Stefano Masier:** Conceptualisation, Writing - review and editing. **Elio Tuci:** Conceptualisation, Methodology, Writing - review and editing, Supervision.

## Acknowledgements

This work was supported by Service Public de Wallonie Recherche under grant n° 2010235 - ARIAC by DIGITAL- WALLONIA4.AI. Computational resources have been provided by the Consortium des Équipements de Calcul Intensif (CÉCI), funded by the Fonds de la Recherche Scientifique de Belgique (F.R.S.-FNRS) under Grant No. 2.5020.11 and by the Walloon Region.

### A Description of the model

In this section, we describe our model following the standard protocol ODD [21] used for individual or agent-based models.

#### A.1 Purpose

The purpose of this model is to study the influence of different sources of information on the behaviour of butterflies over multiple generations. In particular, we study the effect of the cost of acquisition of social or private information on the different genes determining the dispersal behaviour of female butterflies. We also study different types of environment and their impacts on the behaviour.

#### A.2 State variables and scales

The model is composed of two different entities: the agents representing the butterflies and the patches where agents reside.

Agents have three main state variables: an id, a generation and a sex. If the agent is female, it has additional state variables associated with its dispersal behaviour: the probability of dying while dispersing, different genes used for computing the probability of dispersal, two variables bounding the reproductive success of a female and a probability of mutation for the genes. Only females disperse and pass on their genes to the next generation. Patches have an id, an upper limit for their local population and a quality which is the carrying capacity of the patch. See Table A.1 for details.

**Table A.1:** State variables used in the model.

| Parameter | Values/bounds |
| --- | --- |
| Probability of dying while dispersing ($\mu$) | 0.1 |
| Gene enabling the use of social information ($s_1$) | $\{0, 1\}$ |
| Gene enabling the use of private information ($s_2$) | $\{0, 1\}$ |
| Gene weighting social information in the natal patch ($b_1$) | $[-0.2, 0.2]$ |
| Gene weighting social information in the adjacent patch ($b_2$) | $[-0.2, 0.2]$ |
| Gene weighting private information in the natal patch ($a_1$) | $[-0.2, 0.2]$ |
| Gene weighting private information in the adjacent patch ($a_2$) | $[-0.2, 0.2]$ |
| Lower bound for the reproductive success of a female ($r_{\min}$) | $[0.75, 1.0]$ |
| Upper bound for the reproductive success of a female ($r_{\max}$) | 1.0 |
| Probability of mutation of an allele ($P_{\text{mut}}$) | 0.00005 |
| Initial quality of a patch ($K_0$) | 100 |
| Minimum quality of a patch ($K_{\min}$) | 5 |
| Standard deviation of the quality of a patch ($\sigma$) | $\{1; 10; 100\}$ |
| Upper limit for the local population on a patch ($N_{\max}$) | 500 |

All agents in a generation form the population of our model. The environment is composed of a grid of 20×20 patches where edges are linked to form a torus and female agents can disperse to adjacent patches once by generation. A toroidal environment is used to avoid edge effects where agents near the edges would have fewer neighbouring cells, thus simulating an environment that is closer to a natural habitat without fixed boundaries. The model time step corresponds to the lifespan of an agent, and the generations do not overlap. The model is run for 20,000 generations until the evolutionary process arrives at equilibrium. The qualities of the patches vary over time around a mean initial value with a standard deviation. A low standard deviation means that the environment is homogeneous and there is a low evolutionary pressure for the agents to disperse in search of better quality patches; a high standard deviation means that the environment is challenging with patches that can accommodate fewer agents but also patches with higher quality thus reinforcing the evolutionary pressure on the dispersal behaviour.

#### A.3 Process overview and scheduling

A time step in our model corresponds to one generation of agents. Multiple processes happen in the following order:

*Reproduction*: females reproduce and agents for the next generation are born. All agents from the previous generations die.

*Varying quality* : each patch quality changes following the type of environment studied.

*Mating* : newly emerged females mate in each patch before dispersal. If a patch has no males, the females do not mate and will produce no offspring in the next generation.

*Dispersal* : each female estimates its own probability of dispersal and moves (or not) to an adjacent patch.

#### A.4 Design concepts

*Emergence*: the proportion of females dispersing at each generation is directly linked to the genes of the agents influencing the population dynamics.

*Adaptation*: agents adapt to their environment by modifying their genes values accordingly, thus improving their dispersal decision.

*Fitness*: the fitness is modelled implicitly in the reproduction phase by determining the local number of agents for the next generation following the patch quality and the number of females. Females that have made a successful dispersal decision will thus increase the total population size in the next generation.

*Prediction*: agents do not predict future conditions, they only rely on current information to make a dispersal decision.

*Sensing* : individuals can sample the number of neighbours in their local patch and a randomly chosen adjacent patch, as well as the qualities of those patches. This sensing is disabled if the genes allowing the use of information are switched off.

*Interaction*: individuals do not interact with each other, except for mating.

*Stochasticity* : the dispersal decisions, the mortality and the evolution of the genes over time are all stochastic processes related to the agents. The quality of the environment also varies following a normal distribution for all the patches.

*Observation*: population-level data is computed in the simulation and then extracted (e.g. population size). Agent-level data is directly extracted when needed (e.g. genes values for each agent).

#### A.5 Initialisation

All patches are initialised with a quality taken from a normal distribution. 100 agents are initialised on each patch. When initialised, agents have a 50% chance to be male or female. Genes having a continuous value are initialised with a uniform distribution; binary genes have a 50% chance to be 0 or 1.

#### A.6 Input

Environmental variation is introduced by modifying the patches qualities over time, following a normal distribution. The mean of this distribution is equal to 100 and the standard deviation can vary. The higher the standard deviation, the more challenging the environment will be for the agents.

#### A.7 Submodels

Most aspects of our model are directly derived from the one found in [18]. Our main contribution is the addition of two genes enabling the use of social/private information for computing the probability of dispersal, as well as including a weight for modelling reproductive success linked to the acquisition of this information.

*Reproduction*: At each time step, the local number of agents that will spawn in the next generation on a single patch is computed as:

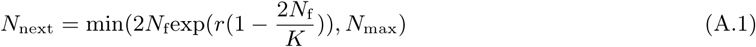

with *N*_f_ the number of females in the previous generation on the patch, *r* = 1 a population growth-rate parameter, *K* the local quality of the patch and *N*_max_ an upper limit for the population. For each new agent, its mother is chosen at random following a weighted distribution with replacement between all the mated females present on the patch. A reproductive weight is computed for each female with:

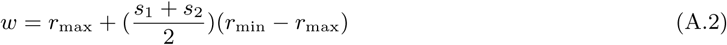

with *r*_min_ and *r*_max_ the bounds of the reproductive weight, *s*_1_ the gene enabling the use of social information and *s*_2_ the gene enabling the use of private information. This equation translates the information cost: the more a female uses information for its dispersal decision, the lower its reproductive success in the local pool of females. This can be viewed as an expense in energy, with females conserving energy having more chances to reproduce. Once a mother is chosen, the new agent inherits directly its genes. Each gene is composed of two alleles (except for *s*_1_ and *s*_2_ that are binary). The sum of these two alleles is used to compute the value of the gene and is bounded (see Table A.1). Each allele has a probability *P*_mut_ to mutate at each generation. When an allele mutates, it is incremented by a value drawn from a Laplacian distribution with mean zero and a standard deviation corresponding to the total interval of the bounds of the gene divided by 40. For binary genes, we simply flip their value. When every new agent has been initialised, all agents from the previous generation die.

*Varying quality* : The quality of a patch varies at each generation following this equation:

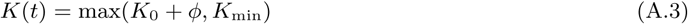

where *ϕ* is an independent normal variable with a mean of zero and a standard deviation *σ*. Quality values are not correlated over time, as we assume the impact of our agents to be non-consequential to the status of the patch compared to unrelated large-scale environmental effects. Dispersal in correlated temporal environments allows for more stable population persistence, while random temporal environments are riskier due to unpredictable and sudden changes in conditions.

*Mating* : If there is a male in their patch, females mate and will be included in the reproduction phase.

*Dispersal* : Females can disperse to an adjacent patch once per generation, and chose to do so following the probability:

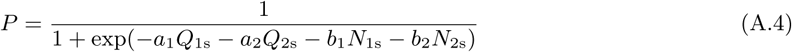

where *a*_1_*, a*_2_*, b*_1_*, b*_2_ are genes related to the dispersal of females. Each female can sample the quality of the natal patch *Q*_1_, the quality of the randomly selected adjacent patch *Q*_2_, the number of individuals in the natal patch *N*_1_ and the number of individuals in the adjacent patch *N*_2_. When a female disperses, she dies with a probability *µ* = 0.1. The use of information is represented by two additional genes *s*_1_ and *s*_2._ Information is enabled with these equations:

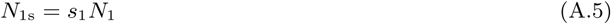

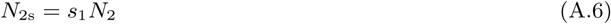

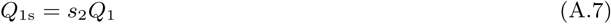

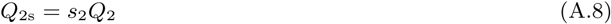

If the value of *s*_1_ or *s*_2_ is equal to 1, the agent will use the associated quantity for its dispersal decision. If it is equal to 0, information will not be used. Females with a restricted range choosing to disperse will move to an adjacent patch at random while females equipped with an extended range will move to the adjacent patch on which they have previously gathered information

